# Control of non-homeostatic feeding in sated mice using associative learning of contextual food cues

**DOI:** 10.1101/247825

**Authors:** Sarah A. Stern, Katherine R. Doerig, Estefania P. Azevedo, Elina Stoffel, Jeffrey M. Friedman

**Author notes:** Correspondence should be addressed to: Jeffrey M. Friedman, Laboratory of Molecular Genetics, Howard Hughes Medical Institute, The Rockefeller University, 1230 York Avenue, New York, NY 10065.

## Abstract

Feeding is a complex motivated behavior controlled by a distributed neural network that processes sensory information to generate adaptive behavioral responses. Accordingly, studies using appetitive Pavlovian conditioning confirm that environmental cues that are associated with food availability can induce feeding even in satiated subjects. However, in mice, appetitive conditioning generally requires intensive training and thus can impede molecular studies that often require large numbers of animals. To address this, we developed and validated a simple and rapid context-induced feeding (ctx-IF) task in which cues associated with food availability can later lead to increased food consumption in sated mice. We show that the associated increase in food consumption is driven by both positive and negative reinforcement and that spaced training is more effective than massed training. Ctx-IF can be completed in ∼1 week and provides an opportunity to study the molecular mechanisms and circuitry underlying non-homeostatic eating. We have used this paradigm to map brain regions that are activated during Ctx-IF with cFos immunohistochemistry and found that the insular cortex, and other regions, are activated following exposure to cues denoting the availability of food. Finally, we show that inhibition of the insular cortex using GABA agonists impairs performance of the task. Our findings provide a novel assay in mice for defining the functional neuroanatomy of appetitive conditioning and identify specific brain regions that are activated during the development of learned behaviors that impact food consumption.

## INTRODUCTION

Associative learning is a fundamental process that enables organisms to make adaptive decisions based on prior outcomes. However, in some cases, associative learning may also lead to maladaptive decisions, as in the case of drug addiction and obesity^1^. Obesity is a growing problem in the developed world, as 30% of adult Americans are currently classified as overweight or obese^2^, ^3^. Though decades of research have elucidated numerous genetic factors underlying obesity, monogenetic disorders do not account for the majority of cases^4^.

Furthermore, environmental factors can also play an important role in overeating and weight gain. Thus, external motivating factors, including easy access to high-energy food, as well as cognitive and emotional cues, can lead to overeating and even binge eating^5–8^. Consistent with this, it has been demonstrated in both humans and rodents that environmental cues that are paired with food delivery can augment subsequent feeding responses. For example, Weingarten used a Pavlovian conditioning paradigm to show that a cue paired with food delivery when rats were hungry could later be used to decrease the latency to eat even when rats were sated^9^. In contrast, an unpaired cue did not affect the latency to eat. More recently, it has been demonstrated in both rats^10^ and mice^11^ that visual and auditory cues can also be used to elicit overeating in sated animals after Pavlovian training.

Pavlovian conditioning is an extremely well-studied paradigm and has been used to study many behaviors, in particular feeding and fear^12^, ^13^. Indeed, Pavlov’s original studies paired food and a sound in dogs^14^, highlighting the powerful motivation of food as a reinforcer. Conditioning experiments can be further subcategorized as either cued or contextual conditioning depending on the type of sensory stimulus that is provided. For example, cued fear conditioning, which measures the freezing response after pairing discrete visual or auditory cues with footshocks, can be demonstrated in as little as one training session^15^. In contrast, conditioning animals to eat even when sated using an analogous cue-induced feeding task with visual or auditory cues requires more extensive training in chronically underfed animals, and it can take as long as one month to fully train an animal^11^.

As an alternative, contextual training has also been used to link associative cues to a response. In previous studies of fear conditioning, a specific context (e.g. a behavioral chamber which provides a variety of different sensory cues) can serve as the conditioned stimulus. For example, it has been shown that placing animals in a novel context paired to a specific stimulus (e.g. footshocks) can induce the subsequent conditioned response (CR) of increased freezing in a single trial^15^. The contextual cues serve as a general predictive environment for a specific outcome rather than the onset of the single sensory input such as an auditory or visual cue as employed in cued Pavlovian conditioning^16^.

While context induced feeding (Ctx-IF) has been used to potentiate food intake in sated rats^17^, this paradigm has not been evaluated in mice. In this report we tested whether context, rather than discrete cues, could be used to induce a conditioned response of overeating in mice. We tested this by placing fasted mice in a novel context where food was made available. We show that, similar to contextual fear conditioning, context can serve as a strong cue to induce overeating in sated mice and the use of this approach leads to similarly robust conditioning when compared to the use of an established cued conditioning protocol with a single defined stimulus. We then used this assay to establish optimal conditions for inducing the conditioned response by varying intertrial intervals. We also defined the emotional valence (positive or negative) driving the feeding response. In order to determine which brain regions might be involved in regulating conditioned overeating, we then used immediate early gene mapping to identify several brain regions that are activated in conditioned animals. Finally, we infused GABA agonists prior to Ctx-IF testing to show that the proper function of the insular cortex, one of the brain regions that is activated by the novel context, is necessary for the conditioned response. The high throughput and convenience of the context induced feeding protocol will facilitate further experiments to define the specific neural populations and circuits that control the overconsumption response.

## METHODS

### Mice

All experiments performed in these studies were approved by the Rockefeller University Institutional Animal Care and Use Committee and were in accordance with the National institute of Health guidelines. Male wild-type C57BL6/J mice (The Jackson Laboratory, Bar Harbor, ME; RRID:IMSR_JAX:000664) were between 10–20 weeks at the start of behavioral testing. Mice were group housed on a 12 h light/dark cycle with *ad libitum* access to water and standard mouse chow, PicoLab Rodent Diet (Lab Diet, St. Louis, MO), except when single housing or fasting is noted below. Mice were handled for 1-2 minutes per day for 4 days prior to the start of behavioral procedures.

### Cued-IF

Mice were trained similarly as described^11^ with the following changes. Mice were food restricted to ∼85% *ad libitum* weight by providing ∼1/3 standard food pellet per day for 1 week prior to the start of training. Weights were taken prior to food restriction and every day before food delivery to ensure adequate weight loss. Following the initial food restriction, mice were trained on a “simple” conditioning paradigm for 5 days and then a “discrimination” phase for 9 days.

Conditioning took place in Coulbourn Instruments (Whitehall, PA) rat-sized operant chambers outfitted with a house light, a blue LED and an automated food well. The house light and blue LED served as the “light” stimuli; the tones consisted of either white noise or a 2.5 kHz tone. During the “simple” conditioning mice were placed into the chamber and the CS+ tone+light combination (balanced between groups) was played for 2-sec every 30-90 sec at random intervals, with a total of 30 cues presented over a 30min conditioning session. Each presentation co-terminated with delivery of a 20 milligram (mg) Precision Pellet (Bioserv, Flemington, NJ, Catalog # F05301). Mice were observed to determine whether they nose-poked to the food well during each cue presentation and the # of pellets consumed was counted at the end of the session. The “discrimination” sessions were conducted in the same manner, except that 15 CS+ (tone+light, paired with food) and 15 CS-(alternative tone+light, not paired with food) presentations were randomly given every 30-90 sec over the course of a 30 min conditioning session. Following the last “discrimination” session mice were returned to *ad libitum* feeding for 5 days before testing. Testing consisted of mice being placed into the test chambers for 2 10-min sessions separated by a 5 min intertrial interval, in which the mouse was placed back into his home cage. In the first session mice were either presented with 10 CS+ presentations or CS-presentations. In the second session mice received 10 presentations of whichever CS they did not receive in the first session. The order of presentation was counterbalanced. In both cases, there were 2 grams (g) of pellets freely available in the food cup. Pellets were weighed after each session to determine consumption. During testing, the investigator was blind to the experimental conditions.

### Context-IF

Prior to habituation mice were given 5-10 chocolate flavored 20mg Precision Pellets during each day of handling to prevent neophobia. Mice were habituated to two different contexts. Contexts were easily distinguishable based on shape, size, floor texture and were in different rooms (Supplementary Figure 1). Habituation consisted of 20 min exposure to each context. Mice were returned to their home cages after habituation, and cages designated as either “Fasted” or “Fed Controls.” One hour later, all cages were changed to prevent food dust on the floor from being consumed, and “Fed controls” continued to have *ad libitum* access to chow while “Fasted” mice did not receive food for the next 24h. The next day mice were trained to associate food delivery with one of the two contexts. Training consisted of 30 min exposure to the context with 2g of Precision Pellets freely available in a food well. Pellets were counted after the training session to determine food consumption. Mice continued to be fasted or fed and were trained again the next day. One hour after the second training session, all mice were returned to *ad libitum* feeding for 48h when testing was conducted. Testing consisted of 20 min exposure to both contexts with 2g of Precision Pellets freely available in a food well. Pellets were again counted after testing to determine food consumption. The order of context presentation during habituation, training and testing sessions were all counterbalanced to prevent ordering effects. During testing, the investigator was blind to the experimental conditions.

**Figure 1.**
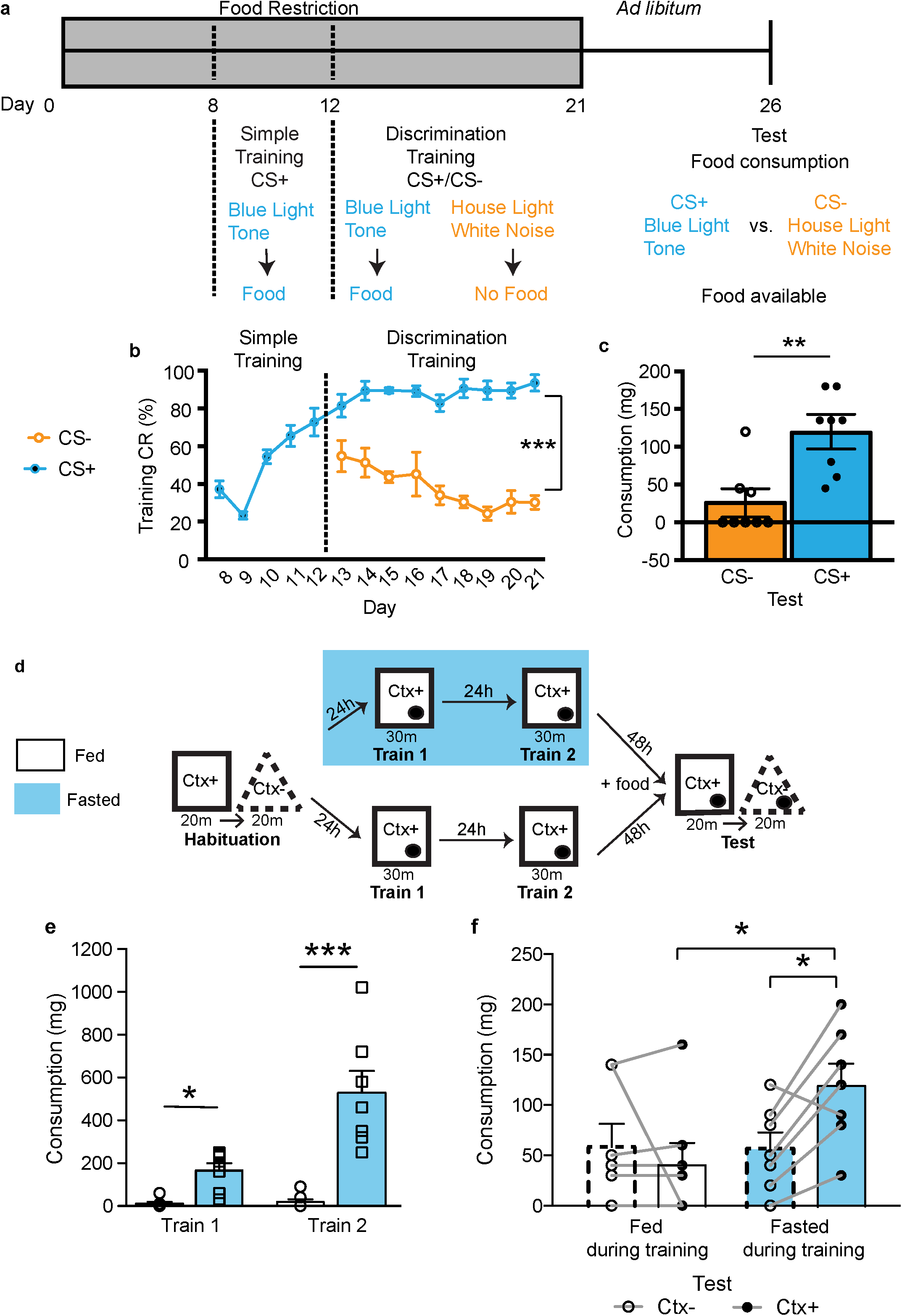
Cue-induced feeding paradigm (Cue-IF) and Context-induced feeding paradigm (Ctx-IF) similarly induce overeating in mice We used an established Pavlovian conditioning paradigm to induce increasing food consumption in sated mice in response to discrete sensory cues (e.g. light and tone) and a novel paradigm utilizing contextual cues to induce a similar response. (**a**) Experimental schedule of Cue-IF. Mice were restricted to 85% body weight (represented by the gray box) from day 0 to day 7 and were conditioned in 2 phases. Phase 1 (Day 8-12) consisted of pairing a tone+light with food delivery (CS+). Phase 2 (Day 13-21) consisted of continuing the original pairing while adding in a second tone+light combination that is not paired with food (CS-). After testing, mice were returned to *ad libitum* feeding and were then tested for their food consumption in response to both the CS+ and CS-. (**b**) Nose pokes into the food well in response to the conditioned stimulus (CS+) compared to nose pokes in response to the unconditioned stimulus (CS-) during training (Days 9-21). The conditioned response (CR) is represented as the percentage (%) of trials in which animals make a nose poke into the food port in response to the CS+ or CS-. (**c**) During testing, the food intake (represented in milligrams, mg) after exposure to CS- and CS + was compared in sated mice, showing a significant increase in response to the conditioned stimulus (CS+). (**a-c**) N=7-8. (**d**) Experimental schedule of Ctx-IF. Mice were habituated to two different novel contexts and then separated into two groups, one fasted overnight (blue) and one that continued to be fed *ad lib*itum (white). They were then trained in one of the two contexts (Ctx+) with food available. After 2 training sessions, all mice were returned to *ad libitum* feeding for 48 hours and were then tested for their food consumption in response to exposure to both the context which had been paired with food during training (Ctx+) and the unpaired context (Ctx−). (**e**) The food intake (represented in milligrams, mg) in response to the paired context (Ctx+) during training, comparing mice that were fasted prior to training (blue bars) and mice that were fed *ad libitum* (white bars). (**f**) During testing, the food intake after exposure to CS- and CS + was compared in both mice that were fasted prior to training and mice that were fed *ad lib*itum. This shows a significant increase in response to the paired context (Ctx+) as compared to the unpaired context (Ctx−), only in mice that were fasted prior to training. (**d-f**) N=7-8. *P<0.05, **P<0.001, ***P<0.001

#### Massed training

The protocol was conducted as described above with the following changes. Training occurred in 2 sessions over the course of one day. Mice were trained for 30 min in the morning, and 5 hours later, were trained again for 30 min. One hour after the second training session, all mice were placed back on *ad libitum* food access until testing occurred 48h later. To increase motivation to consume food during training, a separate group of animals went through the same protocol, but were fasted for 48 hr before the one-day training protocol.

#### Spaced training

The protocol was conducted as described above with the following changes. After the first training session, all mice were placed back onto *ad libitum* food access for 24h. After those 24h, “fasted” mice again had their food removed for the next 24h before the second training session (total of 48 hr between training sessions). One hour after the second training session, all mice were placed back on *ad libitum* food access until testing occurred 48h later.

#### Negative valence

The protocol was conducted as described above with the following changes. Mice were trained as described, but “fasted” mice were further subdivided into two groups. “Positive valence” mice were trained as described above with Precision Pellets freely available during training. “Negative valence” mice were fasted prior to training, but did not receive Precision Pellets in the training context. To control for food intake, they received a paired amount of standard chow in their home cage following training. In order to assess the metabolic status of the mice, mice were weighed during handling and throughout the behavioral protocol.

### Immediate Early Gene Mapping

Mice were trained in Ctx-IF using the original protocol described above, but were tested without food in order to assess activation in response to the context itself. A group of Naïve mice were handled but not trained or tested. Thirty minutes after being tested in Ctx+ or Ctx−, both groups of mice were perfused using 4% paraformaldehyde (PFA). Brains remained overnight in 4% PFA before being transferred to phosphate-buffered saline (PBS) and processed for immunohistochemistry. Brains were sectioned at 50 μm in 4 serial sections. Sections were washed 3x for 15 min in PBST (PBS-0.1% TritonX) and blocked in 3% goat serum in 3% BSA with 0.05% sodium azide. Sections were then incubated overnight with primary cfos antibody (Cell Signaling, Danvers, MA, dilution 1:500), washed 3x with PBST and then incubated for 1 hr at RT with goat anti rabbit AlexaFluor 488 secondary antibody (Thermo Fisher Scientific, Waltham, MA, dilution 1:1000) and washed again 3x in PBST. Sections were mounted using Vectashield with DAPI (Vector Labs, Burlingame, CA) and imaged using an Inverted LSM 780 laser scanning confocal microscope (Zeiss, Thornwood, NY). To ensure valid comparisons, equivalent sections from the Naïve, Ctx− and Ctx+ groups were imaged with the same settings (laser power, gain and digital offset) and any post-acquisition manipulations were also made equivalently in both groups. Quantifications were made using ImageJ thresholding and particle analysis by an experimenter blind to conditions.

### Stereotaxic Surgery and Cannula Implants

Mice were anesthetized with 3.5% isoflurane, placed in a stereotaxic frame (Kopft Instruments, Tujunga, CA), and maintained at 1.5% isoflurane. The craniotomy was performed using a dental drill (Dremel), and stainless steel guide cannulae (28-gauge, Plastics One, Roanoke, VA) were stereotactically implanted to bilaterally target the insular cortex (0.26 mm anterior to the bregma; 3.9 mm lateral from midline; and 3.8 mm ventral from the skull surface) and fixed with dental cement. Mice were monitored for 72h to ensure full recovery, were single housed, and 2-3 weeks later were used in experiments. Fifteen minutes before testing, mice received bilateral injections of muscimol (Sigma, St. Louis, MO) and baclofen (Sigma) (0.05 μg/μL and 0.02 μg/μL, respectively). Injections were delivered in a volume of 0.3 μL per side and used a 33-gauge needle that extended 0.8 mm beyond the tip of the guide cannula. Guide cannulae were connected via polyethylene tubing to Hamilton syringes. All infusions were delivered at a rate of 0.1 μL/min using an infusion pump (Harvard Apparatus, Holliston, MA). The injection needle was left in place for 1 min after the injection to allow complete dispersion of the solution.

To verify proper placement of cannula implants, at the end of the behavioral experiments, mice were perfused with 4% paraformaldehyde in PBS and brains were post-fixed in the same solution. No animals were excluded for improper cannula placement. Forty micrometer coronal sections were cut through the relevant brain region and examined under a light microscope.

### Statistical Analyses

Two-way analysis of variance (ANOVA) followed by the Sidak’s multiple comparisons test, One-way ANOVA followed by Neuman Keul’s multiple comparisons test or unpaired Student’s *t*-test was used for statistical analyses. All graphs represent mean +/- S.E. Statistics were calculated using Graphpad Prism 6. Group sizes were based on power calculations conducted prior to the experiments and no animals were excluded from analysis.

## RESULTS

### Context is sufficient to drive an overeating response in sated mice

In this study we set out to identify and test the function of brain regions that play a role in associative learning to control increased feeding in sated animals. We first confirmed that discrete paired auditory and visual cues can drive overeating as previously described^11^. In these cue induced feeding experiments (Figure 1a), mice were food restricted until they weighed ∼85% of their baseline *ad libitum* weight for 1 week prior to training. Mice were then trained for 5 days to retrieve a food pellet whose delivery was paired to a 2-sec presentation of a conditioned stimulus (CS+), which consisted of a combination of tone + light. Mice were then trained over 9 subsequent sessions to discriminate between the CS+ and an unpaired stimulus, (CS-), balanced between groups (e.g. when blue light or tone was used for CS+, a house light and white noise was used for CS-, and vice versa). We initially evaluated the efficacy of the conditioning by assaying the number of nose pokes into a food well in response to the CS+ vs. CS -. We found that mice responded with nose pokes to the CS+ trials 95% of the time by the last training session, vs. 30% of the time after exposure to the CS- (Two-way ANOVA: CS+ vs CS-: F_(1,112)=_333.2, ***p<0.0001, Session: F_(8,112)_=1.525, p=0.1565, Interaction: F_(8,112)_=2.199, *p=0.0326, followed by Sidak’s multiple comparison test: ***sessions 14-21: CS+ vs. CS-; Figure 1b). In addition, mice retrieved and ate 100% of the delivered pellets after the sixth session indicating that the mice were motivated to eat during training (Supplementary Figure 1). Mice were then fed *ad libitum* for 5 days until their weight returned to baseline after which their food intake post exposure to the CS+ and CS- was tested. The training elicited a robust increase in feeding, as mice ate significantly more during the test in response to the CS+ than to the CS- (t-test, t_(14)_=3.93, **p=0.0015, Figure 1c).

However, from start to finish, this training procedure took a total of 4 weeks using operant chambers, which limits the utility of this approach for generating the larger number of animals that would be required for molecular and functional studies of the neural circuit underlying the observed response. We thus tested whether we could shorten the length of time needed to condition animals by training mice to overeat in response to a specific context rather than in response to discrete cues.

We began by habituating *ad libitum* fed mice to two different novel contexts for one 20 minute session each. The two contexts differed in size, shape, wall color and floor texture. In addition, the two contexts were located in different rooms so that the gross location of the room could not be generalized during testing (Figure 1d, Supplementary Figure 1).

After the habituation session, one group of mice continued to be fed *ad libitum*, while another group of mice were fasted for 24 hours prior to training. During the 30 minute training sessions, mice were placed into one of the contexts (Ctx+) and were able to freely explore and obtain food from a food well. The unpaired context, in which food was not available, was designated as Ctx−. Following each training session mice were returned to their home cages until the next training session. At the conclusion of this two day training period, the mice were fed *ad libitum* in their home cages for two days after which they were tested by returning them, serially into each of the two different contexts (Ctx+ and Ctx−) for twenty minutes in the presence of food. They were again able to freely explore and obtain food from the food well.

A comparison of the food consumption during the training sessions revealed a significant increase in food intake in the mice fasted during training compared to the mice that had been fed during training (i.e; “Fed” controls; two-way ANOVA: Fed vs. Fasted: F_(1,12)_=31.47,***p=0.0001, Test 1 vs. Test 2: F_(1,12)_=14.29, ***p=0.0026, Interaction: : F_(1,12)_=13.21,***p=0.0034, followed by Sidak’s multiple comparison test: ***Train 2: Fed vs. Fasted, ***Fasted: Train 1 vs. Train 2; Figure 1e). As expected, during testing, these Fed control animals ate only a small, similar amount in both Ctx+ (the environment where food was available during training) and Ctx− (the environment where food was not available during training). In contrast, sated mice that had been fasted during training ate significantly more when placed in Ctx+ than in Ctx− during testing (two-way ANOVA: Fed vs. Fasted: F_(1.12)_ = 2.255, p=0.1591, CS: F_(1.12)_ = 2.738, p=0.1239, Interaction: F_(1,12)_=7.797, *p=0.0163, followed by Sidak’s multiple comparison test: *Fasted: CS+ vs. CS-, *CS+: Fasted vs. Fed; Figure 1f). Moreover, the food intake in Ctx+ was similar to that observed in mice responding to the CS+ using the standard cue-induced feeding protocol described previously (see Figure 1a). In addition, the mice that had been fasted during training consumed very little food when placed in Ctx-; indeed, it was equivalent to the amount that Fed control mice consumed in Ctx− (Figure 1f). The observation that the fully trained animals, when refed, failed to increase intake when placed in the Ctx-environment, indicates that the increased intake in Ctx + was not being driven by hunger during the test itself and rather from the learned association of this environment with privation. The order of context in which mice were habituated and tested was counterbalanced to ensure that the order of testing was not a factor in the amount of food consumption. Overall this protocol showed similar efficiency to cue induced Pavlovian feeding despite the fact that the satisfactory completion of a training protocol sufficient to induce an overeating response required a total of 5 days, and just 2 training sessions for ctx-IF compared to an average of 1 month and 14 daily training sessions to condition animals using the cue induced feeding protocol.

### Parameters Influencing Context Induced Feeding

Performance of learning and memory tasks can be affected by even subtle differences in specific aspects of the training paradigm. We thus tested whether varying the length of time between training sessions (e.g. spaced vs. massed training over short periods of times) influenced the robustness of the conditioning, as has been demonstrated for other paradigms^18^. We first tested whether increased food intake after exposure to a specific context could be generated by massed training, in which the training is conducted in two sessions in a single day. Mice were habituated and fasted overnight and were then trained in two sessions with an interval of 5 hours between them (Figure 2a). As expected, the fasted mice consumed more food during training (Figure 2b), but when later were tested in the fed state, they failed to show a difference in food intake between the Ctx+ and the Ctx− (t-test: t_(14)_ =0.097, p=0.925; Figure 2c). We then asked whether the massed protocol would be sufficient to induce overconsumption if there was increased motivation to consume during training. We trained the mice as described above, but after habituation, now fasted them for 48 hours instead of 24 hours (Figure 2d).

**Figure 2.**
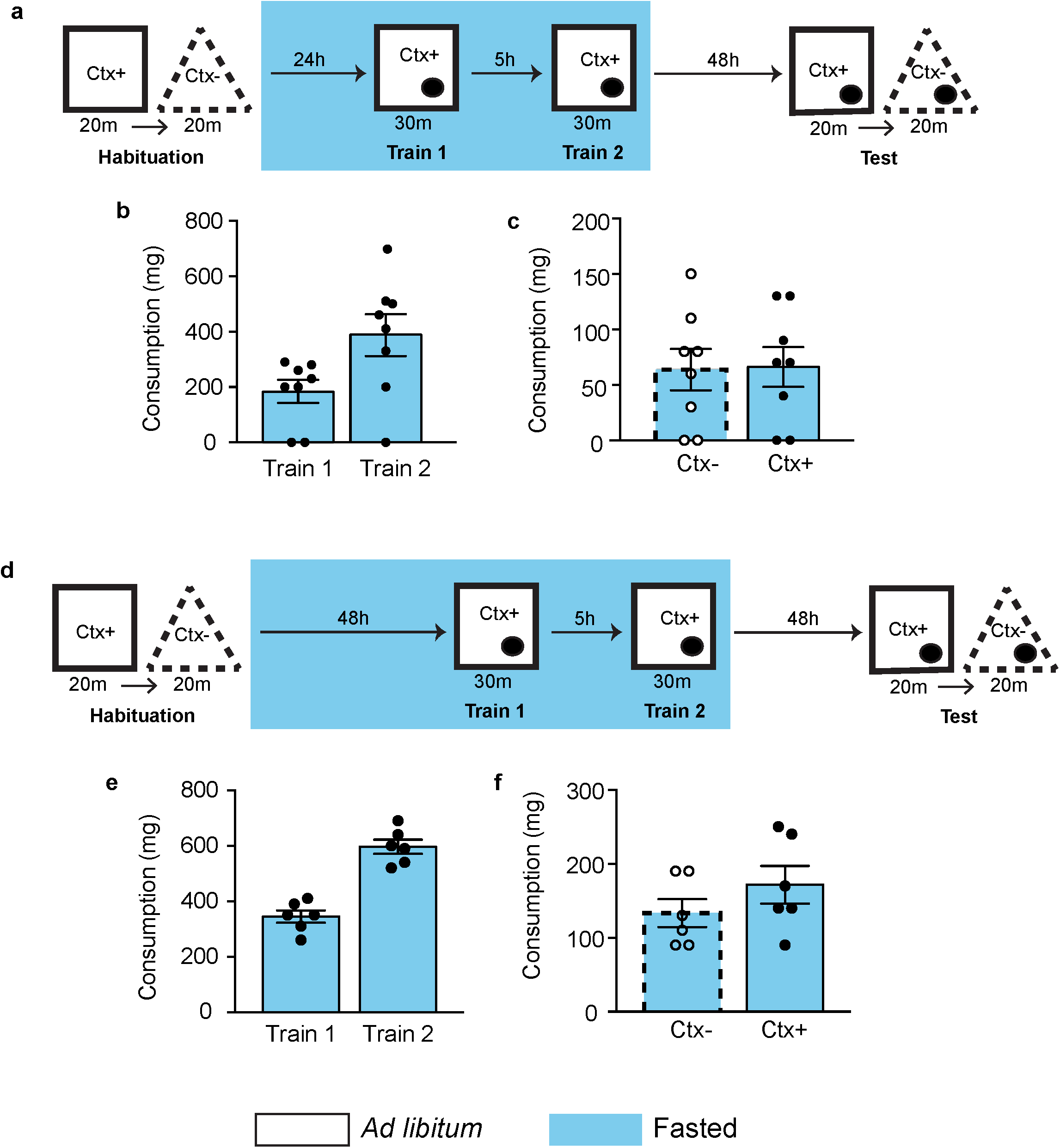
Massed training is not effective to induce context-dependent overconsumption. The impact of massed training trials during Ctx-IF was evaluated. (**a**) Experimental schedule. Mice were habituated and trained as described, but with the 2 training sessions conducted in one day, separated by 5 hours. (**b**) The food intake (represented in milligrams, mg) in response to the paired context (Ctx+) during training in mice that were fasted prior to training. (**c**) During testing, the food intake after exposure to CS− and CS + was compared in mice. There was no increase in response to the paired context (Ctx+) as compared to the unpaired context (Ctx−). (**a-c**) N=8. (**d**) Experimental schedule. Mice were habituated and trained as described, but were fasted for 48 hours prior to the massed training protocol. (**e**) The food intake (represented in milligrams, mg) in response to the paired context (Ctx+) during training in mice that were fasted prior to training. (**f**) During testing, the food intake after exposure to CS- and CS + was compared in mice. There was no increase in response to the paired context (Ctx+) as compared to the unpaired context (Ctx-). (**d-f**) N=6.

Again, we saw that although they are more during training (Figure 2e), the mice failed to show a difference in food intake between Ctx+ and Ctx− during testing (t-test: t_10_=1.208, p=0.2549; Fig 2f) despite the longer period of food restriction and associated increase in motivation. We therefore concluded that massed training conducted over 1 day is not sufficient to induce overeating, regardless of the motivational state of the animal.

We then tested the robustness of the conditioning when the two training sessions were conducted 24 hours apart, or longer. As before, mice were habituated and fasted overnight, after which the first training session in Ctx+ was conducted. The mice were then fed *ad libitum* for 24h after which they were again fasted overnight followed by a second training session in Ctx+. In total, 48 hours separated the two training sessions. Mice were then returned to *ad libitum* food access for two days. The food intake in the sated mice was then compared in Ctx+ vs Ctx− (Figure 3a). During training, fasted mice ate significantly more than fed controls (two-way ANOVA: Training Session: F_(1,6)_=4.5, p=0.0781, Fed vs. Fasted: F_(1,6)_=21.74, **p=0.0035, Interaction: F_(1,6)_=4.5, p=0.0781; Figure 3b). During testing, mice who had been fasted during training consumed significantly more food in response to the training context, Ctx+, than to the Ctx− (two-way ANOVA: Ctx: F_(1,6)_=29.99, **p= 0.0015; Fasted vs Fed: F_(1,6)_=10.22, *p=0.0187;Interaction: F_(1,6)_=11.02, *p= 0.0160, followed by Sidak’s multiple comparison test: **CS+ - fasted vs. fed, **fasted – CS+ vs. CS-; Figure 3c). As before, mice that had been fed during the training sessions ate only a small amount of food, in similar amounts, in the two environments, Ctx+ and Ctx-. These data show that massed training in a single day is not effective, while conducting the training trials over an interval of at least 48 hours generates a robust association between the training Ctx+ and food intake in sated mice.

**Fig. 3.**
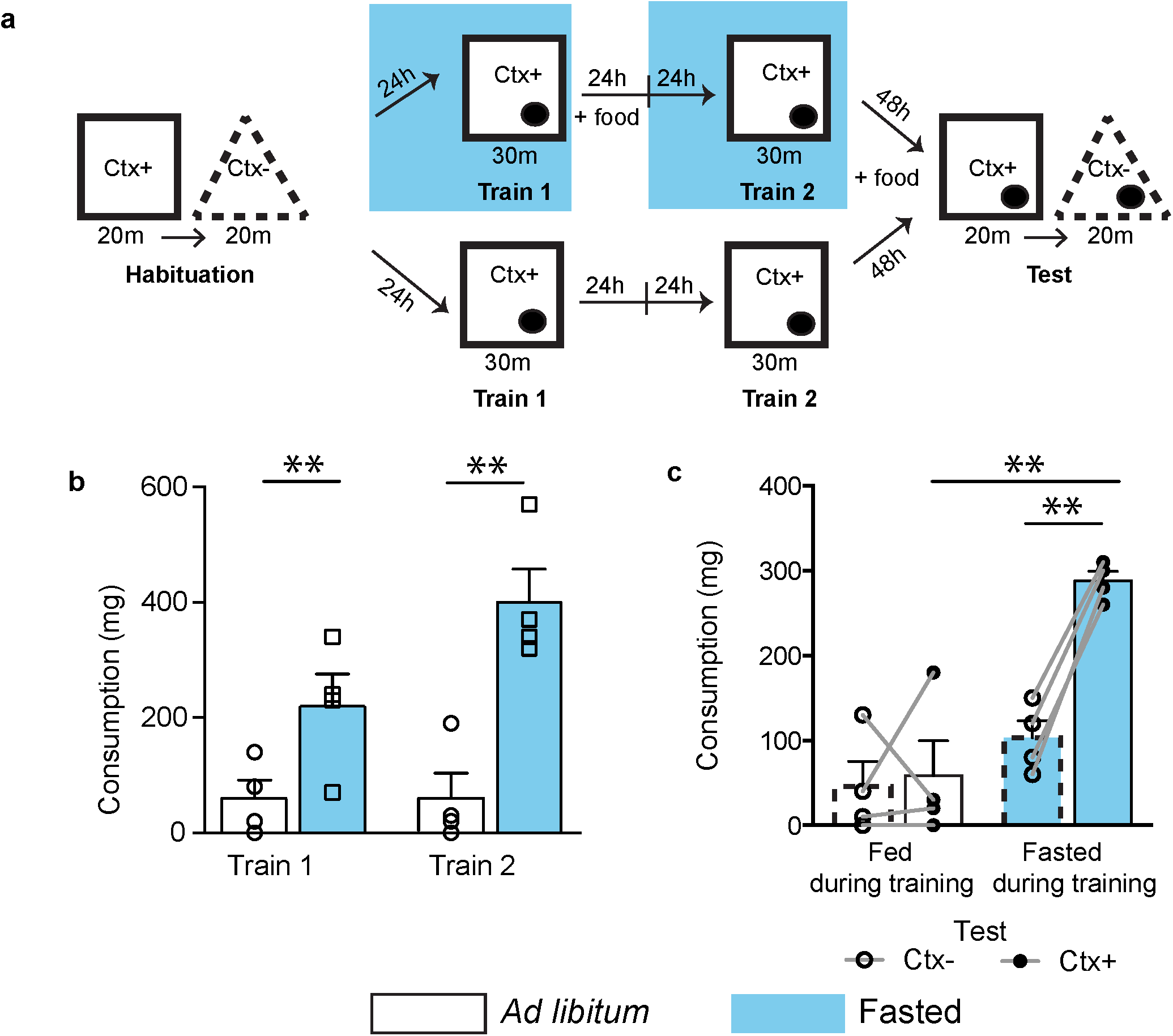
Spaced training reliably induces context-dependent overconsumption. **(a)**Experimental schedule. Mice were habituated and trained as described, but Fasted mice (top) were trained with 2 sessions, separated by 48h and were given *ad libitum* access for 24h after training session 1. Fed controls (bottom) were trained at the same time without any prior food deprivation. (**b**) The food intake (represented in milligrams, mg) in response to the paired context (Ctx+) during training, comparing mice that were fasted prior to training sessions (blue bars) and mice that were fed *ad libitum* (white bars). (**c**) During testing, the food intake after exposure to Ctx− and Ctx+ was compared in both mice that were fasted prior to training and mice that were fed *ad libitum*. This shows a significant increase in response to the paired context (Ctx+) as compared to the unpaired context (Ctx-), only in mice that were fasted prior to training. N=4. **P<0.01, ***P<0.001

### Context-Induced Feeding has components of negative and positive valence

We next asked whether the increased food intake in the conditioned animals’ response was driven primarily by negative valence (e.g. hunger) or positive valence (e.g. food reward). We hypothesized that if the conditioned behavior (food intake) was driven primarily by food reward (and not by hunger), mice that were fasted and then trained in the Ctx+ without receiving food there (negative valence group, Figure 4a, middle) would not show potentiated feeding during testing sessions. In contrast, if the response was driven primarily by hunger, the negative valence group would still show a potentiated feeding response in a subsequent testing session, similar to mice that were given food during the training sessions in Ctx+ (positive valence group, Figure 4a, top).

**Fig. 4.**
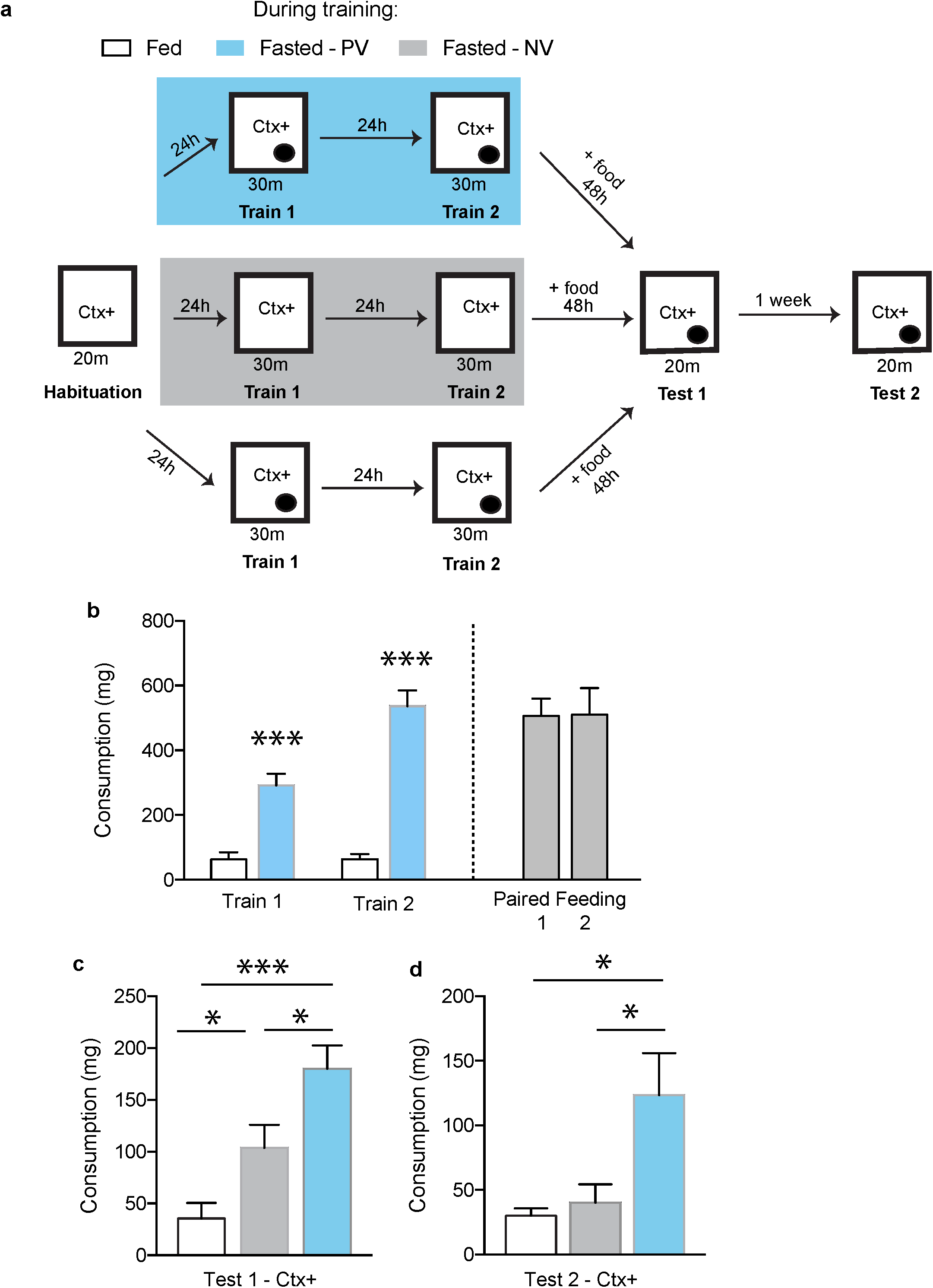
Ctx-IF is driven primarily by positive reinforcement. The role of hunger vs food reward to drive increased consumption during Ctx-IF testing was tested. (**a**) Experimental schedule. Positive valence mice (top, Blue) were fasted prior to training and trained with food available as described, whereas negative valence mice (middle, Grey) were fasted prior to training, but trained without food availability. Fed controls (bottom) were trained with food availability, but were not fasted prior to training, as described. (**b**) The food intake (represented in milligrams, mg) in response to the paired context (Ctx+) during training, comparing mice that were fasted prior to training sessions (Positive valence mice, blue) and mice that were fed *ad libitum* (Fed controls, white). Paired amounts of food were given to the negative valence group (gray). (**c**) During Test 1, the food intake after exposure to Ctx− and Ctx+ was compared in positive valence mice, negative valence mice and fed controls. Positive valence mice mice consume significantly more than fed controls and negative valence mice in response to Ctx+ during testing. (**d**) During Test 2 the food intake after exposure to Ctx− and Ctx+ was compared in positive valence mice, negative valence mice and fed controls. Positive valence mice mice consume significantly more than fed controls and negative valence mice in response to Ctx+ during testing. N=7-8. *P<0.05, **P<0.01, ***P<0.001.

Body weight did not differ between groups before training or during habituation (Supplementary Figure 2). As expected, body weight of both the positive and negative valence groups, which were fasted, dropped significantly during the training protocol. During training, fed controls (Figure 4a, bottom) ate very little, while the positive valence group, which is trained with food available in the training context, ate significantly more during training, as they did in previous experiments (Two-way ANOVA: ***Train 1 vs 2: F_(1,23)_= 34.86, P<0.0001, ***Fed vs.

Fasted: F_(1,23)_= 49.31, P<0.0001, ***Interaction: F_(1,23)_= 34.86, followed by Sidak’s multiple comparison: ***Train 1: fed vs. positive, ***Train 2: fed vs. positive; Figure 4b). The negative valence group was trained without food and therefore no consumption measurement was made during training sessions. However, the mice were then given amounts of food paired to that of the fasted group during the intertrial intervals to ensure that their metabolic status was equivalent to the positive control group (Figure 4b, right). Body weight of both the positive and negative valence groups returned to normal within the first 24h after training and remained comparable to the fed control group by the time of the first testing session (Supplementary Figure 2). As expected, fed controls consumed only a small amount of food in Ctx+ during the testing session. In contrast, the positive valence group that was fasted prior to training and then given access to food in the Ctx+ during training, ate significantly more in the Ctx+ than the fed controls during testing (Figure 4c), as in previous experiments (see Figures 2 and 3).

Interestingly, mice in the negative valence group exhibited an intermediate effect. While they ate significantly more than the fed controls in response to the Ctx+, they consumed significantly less food than the mice in the positive valence group (one-way ANOVA; F_(2,36)_=11.34, ***p=0.0002, followed by Neuman Keul’s multiple comparison test: ***Fed vs. Positive, *Positive vs. Negative; *Fed vs. Negative; Figure 4c). Furthermore, when we tested these animals again 1 week later, mice in the positive valence group still ate significantly more than both the fed controls and the negative valence group, but mice in the negative valence group no longer consumed more than the fed controls (one-way ANOVA: F_(2,13)_=4.891, *p=0.0261, followed by Neuman Keul’s multiple comparison test: *Fed vs. Positive, *Positive vs. Negative; Figure 4d), indicating that the memory that drives overconsumption is both driven by reward and is long-lasting. We therefore concluded that, although negative valence contributes to the potentiated feeding response, the maximal response is driven by the reward value of food received during training sessions and the effect of positive valence is more durable.

### Mapping of Brain Regions Activated by Context Induced Feeding

We next determined which brain regions are activated by Ctx+ in conditioned animals using immediate early gene mapping with cFos as a marker for neural activation^19^. As described above, fasted mice were trained to associate food with a specific context, then refed and euthanized 30 minutes after placement in Ctx+. Another group of naïve mice that were not trained, but were euthanized at the same time as the Ctx+ group, served as controls and a third group was exposed to the unpaired context (Ctx-) before being euthanized (Figure 5a). We then performed immunohistochemistry using cFos to compare its levels in Ctx+ mice to Ctx-mice and Naïve controls. In this analysis we focused on higher-order brain regions because of the known functional relationship of several cortical and subcortical structures with learning and memory. We found that subpopulations of neurons in the following brain regions are activated after exposure to the Ctx+ compared to the control groups, including the insular cortex (IC), central amygdala, lateral septum and lateral hypothalamus (t-test, insular cortex: t_2_=2.4, *p=0.0221, central amygdala: t_2_=5.277, *p=0.0341, lateral septum: t_2_=6.656, *p=0.0218, lateral hypothalamus: t_2_=7.992, *p=0,0153, Figure 5b). Each of these brain regions has been previously shown to play a role in the control of food intake^20–28^. Conversely, In the paraventricular thalamus, we observed increased cFos in Ctx-compared to Ctx+ (t-test: t_2_=2.854, *p=0.0462, Supplementary Figure 3). In the basolateral amygdala we found significant, but equivalent, cFos staining in both the Ctx+ and Ctx− (t-test: t_2_=0.1153, p=0.9918, while a number of regions showed very little cFos in any condition, including S1 and the caudate putamen (Supplementary Figure 3). Although naïve mice displayed almost no cfos staining in almost all of these regions, they did exhibit prominent cfos staining in other regions such as the dentate gyrus (Supplementary Figure 4).

**Fig. 5.**
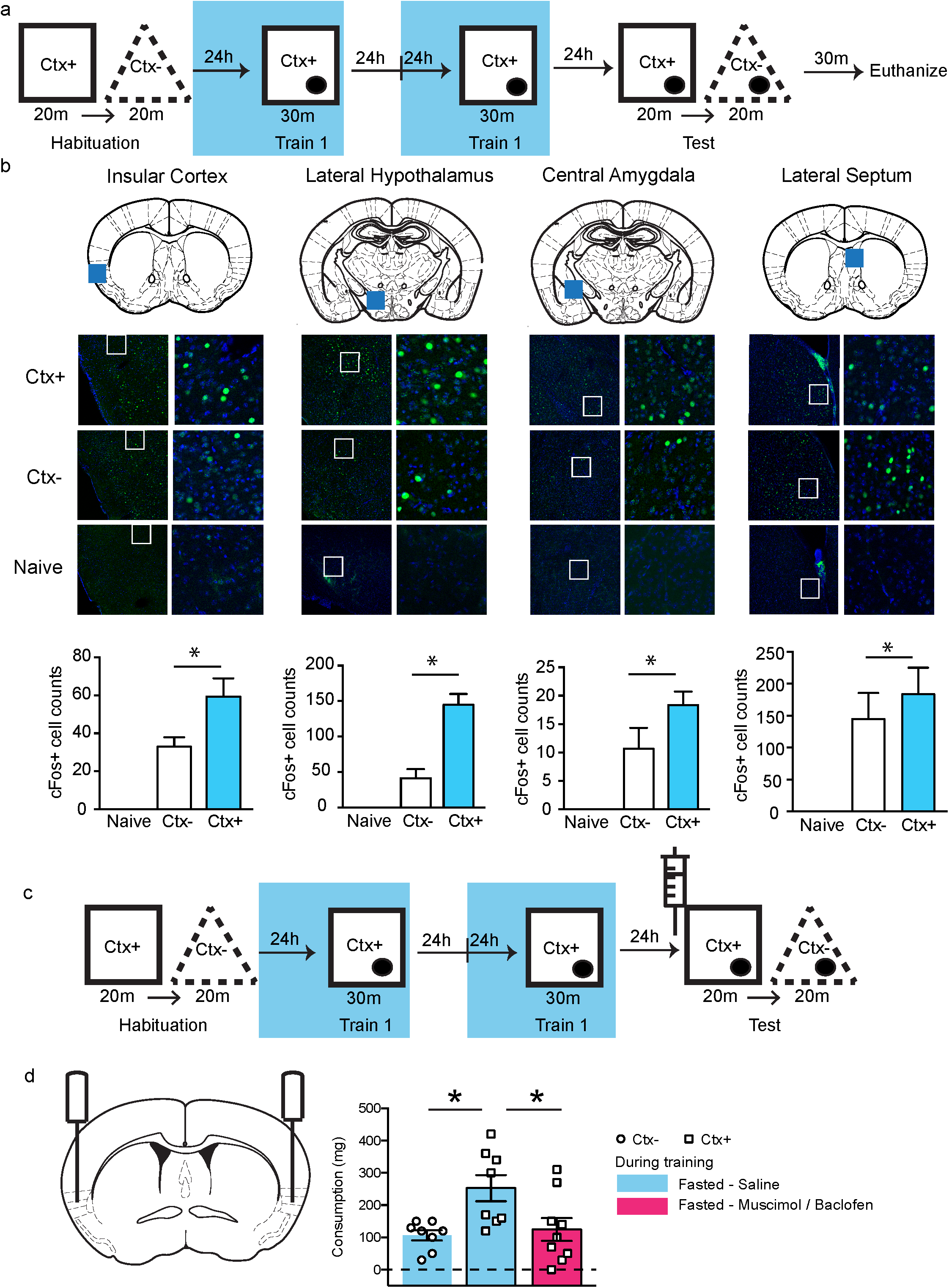
CIF activates the insular cortex, which is required for overconsumption The anatomic loci activated in response to the paired context (Ctx+) and unpaired contex (Ctx-) were mapped. (**a**) Experimental schedule. Mice were habituated and trained as described, and were then euthanized 30 minutes following Ctx-IF training. (**b**) Increased activation as measured by cFos (green) in the insular cortex, lateral septum, central amygdala and lateral hypothalamus in mice, following ctx-IF testing in Ctx+ (top), compared to Ctx− (middle) and Naïve mice (bottoms). DAPI is stained in blue. White inset boxes are magnified to the right. N=3. (**c**) Experimental schedule. Mice were habituated and trained as described, but were injected with saline or muscimol/baclofen solution 15 minutes before Ctx-IF testing into the insular cortex. (**d**) During testing, food intake was compared in Fasted mice who were either injected with saline (Blue) or muscimol/baclofen (Pink) prior to the testing session. Saline-injected mice consume significantly more in response to Ctx+ than Ctx-. In contrast, fasted mice injected with muscimol/baclofen (Pink) eat significantly less in the Ctx+ than saline injected mice. N=8-9. *P<0.05.

We found the insular cortex to be of particular interest because a previous study suggested that the insula projects to multiple efferent sites that play a role in feeding^29^. A detailed analysis of the cFos staining throughout the insular cortex revealed that although the number of cFos+ cells was significantly increased in the Ctx+ group compared to Ctx-overall, post-hoc comparisons did not reveal any particular regions of the insular cortex to be significantly enriched (Supplementary Figure 5). We therefore chose to inactivate the insular cortex in approximately the middle of its A/P axis (+0.6 A/P from bregma) and test its role in Ctx-IF, using the GABA agonists, muscimol and baclofen. Mice were trained as described to exhibit Ctx-IF and 20 minutes before testing were injected with either saline or a combination of muscimol and baclofen, directly into the IC (Figure 5c). We found that mice injected with saline ate more in the Ctx+ vs. the Ctx-environment, as expected. In contrast, mice in which the insular cortex was inhibited just prior to training did not overeat and instead consumed the same amount of food in Ctx+ as saline-injected mice did in Ctx-(one-way ANOVA, F(1,6)=21.74, **p=0.0098, followed by Neuman Keul’s multiple comparison Test: *Saline-Ctx1 vs. Saline Ctx+; *Saline-Ctx+ vs. Muscimol/Baclofen-Ctx+; Figure 5d). Overall these data suggest that neurons in the insular cortex are activated when animals are trained to associate cues with food availability and that proper functioning of this region is required for the development of this association.

## DISCUSSION

Feeding is a complex motivated behavior and in recent years numerous neural circuits that regulate the homeostatic control of ingestion have been identified^30^, ^31^. However, feeding can also be influenced by external cues that predict food availability and it is well established that Pavlovian conditioning of animals can lead to increased food intake even in fed, satiated animals^9–11^. However, in contrast to the substantial progress that has been made in understanding the neural mechanisms that control homeostatic feeding, our understanding of the mechanisms that control cue-induced feeding is limited. The importance of delineating the mechanisms responsible for this behavior is important because cue-induced overeating has been demonstrated repeatedly in overweight and obese human subjects and may also underlie aspects of compulsive eating and binge-eating disorder^32^, ^33^. We thus set out to develop a robust assay for eliciting behavioral conditioning of feeding in mice with the aim of identifying neural sites that play a role in the genesis of this behavior. Here we report the validation of a rapid and robust protocol for context-induced feeding in mice and its use to identify brain regions that are activated by the conditioning stimulus, which is a specific context. We also provide evidence that normal functioning of the insular cortex is required for the development of this response.

Current paradigms in use as models of overeating are of limited utility for studying associative mechanisms that convey food reward and lead to increased food consumption. For example, diet induced obese mice (DIO) only have access to a high fat diet (HFD)^34^, and thus learning is not tested in this model. Other studies have examined the mechanisms underlying cue-induced instrumental responses, that is, the act of obtaining a food reward predicted by a specific sensory cue^35^, ^36^. However, these studies typically examine mice that are food-restricted and are therefore highly motivated to seek a food reward. Thus, because these mice are only tested when hungry, it is difficult to disentangle the contribution of homeostatic feeding in these assays.

As an alternative, several groups have shown that cue induced feeding can be used to study increased consumption in sated subjects. Unfortunately, also shown here, the task requires extensive training over a long period of time, thus limiting its utility for using optogenetics or equivalent methods to dissect the neural mechanisms that control this set of responses. This is because optogenetic and chemogenetic methods to modulate neural activity require that one perform stereotaxic surgeries (∼1-2 week for recovery) with viral injections (∼2-4 weeks for viral expression) further extending the time required to complete a study. In addition, the limited throughput using operant chambers in a cue induced feeding protocol also makes it difficult to generate a sufficiently large cohort of animals. We therefore set out to employ a version of this task that offered greater throughput and less extensive training with which to study the mechanisms and circuitry underlying conditioned overeating. We reasoned that since context is associated with defined sensory inputs, it would be possible to assign motivational salience to a specific environment (referred to as Ctx+) rather than the use of discrete cues. The utility of context as an associative stimulus for feeding in sated mice was validated even when using a protocol that employed few as 2 training sessions over 2 days.Shorter training periods however were not effective for inducing food intake in fed mice. That spaced training is more effective than massed training is consistent with the vast majority of learned behaviors^37–39^. Thus, we showed that two training sessions conducted 5 hours apart was not sufficient to induce increased consumption during the Ctx-IF test. The failure of the trained mice to eat more in the testing session did not appear to be due to decreased hunger during the second training session, as there was no statistical difference between food consumption during the two training sessions. Indeed, in all the experiments, mice in fact tended to eat more during the second training session, indicating that they have already begun to learn the significance of the context by the second session.

Food intake can be driven either by the relief of negative valence (ie. the unpleasant experience of hunger) or by positive valence which refers to the reinforcing or reward value of food^40^,^41^. We thus tested whether the increased intake of food in sated mice after training was driven by positive or negative valence. We found that, in ctx-IF, the maximal consumption response is driven primarily by reward, as mice given food during training (positive valence) eat significantly more in response to the Ctx+ compared to Fed controls and to mice trained without the food reward (negative valence). Recent studies have suggested that activation of AGRP neurons in the hypothalamus increases feeding by conveying negative valence (that is alleviated by feeding)^42^ and can indirectly induce feeding via indirect projections to the insular cortex, which we show contributes to this behavior^43^. Because we saw that the negative valence group eats an intermediate amount of food, these results raise the possibility that mice overeat in response to the Ctx+ in part because the context reminds them of the unpleasant sensation of hunger that was previously experienced in that environment. However, because this group did not eat as much as animals who received food during training, this suggests that the maximal increased feeding response is also driven by the pleasure derived from the food received there (positive valence).

What then are the neural mechanisms underlying non-homeostatic eating? In this study we used cFos as a marker of neural activity to determine which brain regions are activated following ctx-IF. We found that a number of cortical and subcortical brain regions including insular cortex, central amygdala, lateral septum and lateral hypothalamus were activated in sated animals placed into the Ctx+. These findings are consistent with a previous study that measured Fos induction during and after the training portion of a cue-IF^44^ paradigm. This paper also reported that Fos induction differed significantly between early and late training sessions in a number of forebrain regions including infralimbic and prelimbic, insular cortex, dorsomedial striatum, and hypothalamus (lateral and paraventricular) in tone-food paired groups. In contrast, we performed an unbiased screen of brain regions activated by context in sated mice after training had been completed and the association had been fully established. The brain regions we identified in this manner partially overlapped with those identified during training including the insular cortex, as well as the amygdala and lateral septum. We also found activation of the lateral hypothalamus (LH), which is consistent with the finding that the mice consume more food. This finding is also consistent with neural tracing studies using pseudo-rabies virus (PRV) to map brain regions synaptically connected to peripheral areas controlling motor and autonomic aspects of feeding^29^. The insular cortex was one of a small number of brain regions that were common to the descending neural circuits projecting to all of the peripheral sites, indicating that it is likely involved in an integrative function in generating feeding behavior^29^.Although some studies suggest different functional roles for rostral and caudal insular cortex, we did not see any subregional differences when we examined cFos levels after Ctx-IF.

While cFos is a reliable marker for neural activation, it does not provide any insight into the specific neural populations involved in generating the behavior. Recently, PhosphoTRAP, a method that can be used to profile neurons in activated by a specific stimulus, has been developed, thus making it possible to identify the specific neural subpopulations that are activated by the conditioned stimulus (i.e; context)^45^. The identification of markers for the activated neurons in the aforementioned brain regions will enable testing of their function and the ways in which these higher order circuits interact with brain regions that control homeostatic feeding. Another recently developed immuno-precipitation method can also be used to determine precisely the neural populations that connect to these homeostatic centers^46^. Both of these methods are currently being used to define the neural populations that are specifically activated by context in trained animals and that project to homeostatic centers.

Finally, we also used GABA agonists, muscimol and baclofen, to test whether the insular cortex is necessary for the development of ctx-IF. We chose to focus on the insular cortex because of the observation that it indirectly projects to multiple efferent sites that are activated during feeding. In addition, the IC receives projections from the basolateral amygdala, which is critical for associative learning and the IC also sends projections to the central amygdala, lateral hypothalamus and nucleus accumbens, brain regions that are also known to impact feeding behavior^47–50^. Furthermore, other studies have also shown that the insular cortex is required for learning cue-reward associations^43^,^51^. However, a direct role in overconsumption for the IC has not yet been demonstrated and other studies using GABA agonists have failed to show a role for the insular cortex to control homeostatic feeding^43^,^52^.Thus our results suggest a role for the insular cortex in identifying cues that predict food availability potentially accounting for binge-like episodes in response to those cues.

In summary, we have developed a context induced feeding assay that can be rapidly and efficiently employed to condition mice to eat more in response to specific environmental cues. We used this task to study brain circuits that are activated during conditioning and further provide evidence for a functional role of the insular cortex. These findings will provide the basis for future experiments to identify the specific neural populations and establish their function as well as the neural connections through which they exert this effect.

## Acknowledgements

This work was supported by grants from the US National Institute of Health (F32-DK107077 to S.A.S) and the JBC Foundation to J.M.F. The study was conceived and designed by S.A.S and J.M.F. Experiments were conducted by S.A.S, K.R.D. EP.A and E.S. The manuscript was written by S.A.S and J.M.F. with comments from all authors. The authors declare no competing financial interests. We thank the staff of the CBC at the Rockefeller University for technical support.

## REFERENCES

1. Volkow ND, Wise RA. How can drug addiction help us understand obesity? Nat Neurosci 2005; 8: 555–560.

2. Flegal KM, Carroll MD, Kit BK, Ogden CL. Prevalence of obesity and trends in the distribution of body mass index among US adults, 1999-2010. JAMA 2012; 307: 491–497.

3. Ogden CL, Yanovski SZ, Carroll MD, Flegal KM. The epidemiology of obesity. Gastroenterology 2007; 132: 2087–2102.

4. Farooqi IS, O’Rahilly S. Monogenic obesity in humans. Annu Rev Med 2005; 56: 443–458.

5. Zheng H, Berthoud H-R. Neural systems controlling the drive to eat: mind versus metabolism. Physiology 2008; 23: 75–83.

6. Berridge KC, Ho C-Y, Richard JM, DiFeliceantonio AG. The tempted brain eats: pleasure and desire circuits in obesity and eating disorders. Brain Res 2010; 1350: 43–64.

7. Ziauddeen H, Alonso-Alonso M, Hill JO, Kelley M, Khan NA. Obesity and the neurocognitive basis of food reward and the control of intake. Adv Nutr 2015; 6: 474–486.

8. Hill JO, Peters JC. Environmental contributions to the obesity epidemic. Science 1998; 280: 1371–1374.

9. Weingarten HP. Conditioned cues elicit feeding in sated rats: a role for learning in meal initiation. Science 1983; 220: 431–433.

10. Petrovich GD. Forebrain circuits and control of feeding by learned cues. Neurobiol Learn Mem 2011; 95: 152–158.

11. Walker AK, Ibia IE, Zigman JM. Physiology & Behavior Disruption of cue-potentiated feeding in mice with blocked ghrelin signaling. Physiol Behav 2012; 108: 34–43.

12. Maren S. Neurobiology of Pavlovian fear conditioning. Annu Rev Neurosci 2001; 24: 897–931.

13. Everitt BJ, Cardinal RN, Parkinson JA, Robbins TW. Appetitive Behavior. Ann N Y Acad Sci 2003; 985: 233–250.

14. Pavlov IP. Lectures on conditioned reflexes (WH Gantt, Trans.). New York: International 1928;

15. Stern SA, Kohtz AS, Pollonini G, Alberini CM. Enhancement of memories by systemic administration of insulin-like growth factor II. Neuropsychopharmacology 2014; 39: 2179–2190.

16. Phillips RG, LeDoux JE. Differential contribution of amygdala and hippocampus to cued and contextual fear conditioning. Behav Neurosci 1992; 106: 274–285.

17. Petrovich GD, Ross C a., Gallagher M, Holland PC. Learned contextual cue potentiates eating in rats. Physiol Behav 2007; 90: 362–367.

18. Carew TJ, Pinsker HM, Kandel ER. Long-term habituation of a defensive withdrawal reflex in aplysia. Science 1972; 175: 451–454.

19. Sagar SM, Sharp FR, Curran T. Expression of c-fos protein in brain: metabolic mapping at the cellular level. Science 1988; 240: 1328–1331.

20. Stuber GD, Wise RA. Lateral hypothalamic circuits for feeding and reward. Nat Neurosci 2016; 19: 198–205.

21. Frank S, Kullmann S, Veit R. Food related processes in the insular cortex. Front Hum Neurosci 2013; 7: 499.

22. Sweeney P, Yang Y. An excitatory ventral hippocampus to lateral septum circuit that suppresses feeding. Nat Commun 2015; 6: 10188.

23. Scopinho AA, Resstel LBM, Corrêa FMA. alpha(1)-Adrenoceptors in the lateral septal area modulate food intake behaviour in rats. Br J Pharmacol 2008; 155: 752–756.

24. Ono T, Nishino H, Sasaki K, Fukuda M, Muramoto K-I. Role of the lateral hypothalamus and the amygdala in feeding behavior. Brain Res Bull 1980; 5: 143–149.

25. Zhang Q, Li H, Guo F. Amygdala, an important regulator for food intake. Front Biol 2011; 6: 82–85.

26. Cai H, Haubensak W, Anthony TE, Anderson DJ. Central amygdala PKC-δ(+) neurons mediate the influence of multiple anorexigenic signals. Nat Neurosci 2014; 17: 1240–1248.

27. Douglass AM, Kucukdereli H, Ponserre M, Markovic M, Gründemann J, Strobel C et al. Central amygdala circuits modulate food consumption through a positive-valence mechanism. Nat Neurosci 2017; 20: 1384–1394.

28. Sweeney P, Yang Y. An Inhibitory Septum to Lateral Hypothalamus Circuit That Suppresses Feeding. J Neurosci 2016; 36: 11185–11195.

29. Pérez CA, Stanley SA, Wysocki RW, Havranova J, Ahrens-Nicklas R, Onyimba F et al. Molecular annotation of integrative feeding neural circuits. Cell Metab 2011; 13: 222–232.

30. Morton GJ, Cummings DE, Baskin DG, Barsh GS, Schwartz MW. Central nervous system control of food intake and body weight. Nature 2006; 443: 289–295.

31. Gao Q, Horvath TL. Neurobiology of feeding and energy expenditure. Annu Rev Neurosci 2007; 30: 367–398.

32. Jansen A, Havermans RC, Nederkoorn C. Cued Overeating. In: Handbook of Behavior, Food and Nutrition. Springer, New York, NY; 2011. p. 1431–1443.

33. Wardle J. Conditioning processes and cue exposure in the modification of excessive eating. Addict Behav 1990; 15: 387–393.

34. Wang C-Y, Liao JK. A mouse model of diet-induced obesity and insulin resistance. Methods Mol Biol 2012; 821: 421–433.

35. Yager LM, Robinson TE. Cue-induced reinstatement of food seeking in rats that differ in their propensity to attribute incentive salience to food cues. Behav Brain Res 2010; 214: 30–34.

36. Morris MJ, Beilharz JE, Maniam J, Reichelt AC, Westbrook RF. Why is obesity such a problem in the 21st century? The intersection of palatable food, cues and reward pathways, stress, and cognition. Neurosci Biobehav Rev 2015; 58: 36–45.

37. Commins S, Cunningham L, Harvey D, Walsh D. Massed but not spaced training impairs spatial memory. Behav Brain Res 2003; 139: 215–223.

38. Kogan JH, Frankland PW, Blendy JA, Coblentz J, Marowitz Z, Schütz G et al. Spaced training induces normal long-term memory in CREB mutant mice. Curr Biol 1997; 7: 1–11.

39. Wingard JC, Goodman J, Leong K-C, Packard MG. Differential effects of massed and spaced training on place and response learning: A memory systems perspective. Behav Processes 2015; 118: 85–89.

40. Berridge KC. Motivation concepts in behavioral neuroscience. Physiol Behav 2004; 81: 179–209.

41. Sternson SM, Nicholas Betley J, Cao ZFH. Neural circuits and motivational processes for hunger. Curr Opin Neurobiol 2013; 23: 353–360.

42. Betley JN, Xu S, Cao ZFH, Gong R, Magnus CJ, Yu Y et al. Neurons for hunger and thirst transmit a negative-valence teaching signal. Nature 2015; 521: 180–185.

43. Livneh Y, Ramesh RN, Burgess CR, Levandowski KM, Madara JC, Fenselau H et al. Homeostatic circuits selectively gate food cue responses in insular cortex. Nature 2017; 546: 611–616.

44. Cole S, Hobin MP, Petrovich GD. Appetitive associative learning recruits a distinct network with cortical, striatal, and hypothalamic regions. Neuroscience 2015; 286: 187–202.

45. Knight Z a., Tan K, Birsoy K, Schmidt S, Garrison JL, Wysocki RW et al. Molecular profiling of activated neurons by phosphorylated ribosome capture. Cell 2012; 151: 1126–1137.

46. Ekstrand MI, Nectow AR, Knight Z a., Latcha KN, Pomeranz LE, Friedman JM. Molecular profiling of neurons based on connectivity. Cell 2014; 157: 1230–1242.

47. Reynolds SM, Zahm DS. Specificity in the projections of prefrontal and insular cortex to ventral striatopallidum and the extended amygdala. J Neurosci 2005; 25: 11757–11767.

48. Saper CB. Convergence of autonomic and limbic connections in the insular cortex of the rat. J Comp Neurol 1982; 210: 163–173.

49. Yasui Y, Breder CD, Saper CB, Cechetto DF. Autonomic responses and efferent pathways from the insular cortex in the rat. J Comp Neurol 1991; 303: 355–374.

50. Allen GV, Saper CB, Hurley KM, Cechetto DF. Organization of visceral and limbic connections in the insular cortex of the rat. J Comp Neurol 1991; 311: 1–16.

51. Kusumoto-yoshida I, Liu H, Chen BT, Fontanini A, Bonci A. Central role for the insular cortex in mediating conditioned responses to anticipatory cues. 2014;Available from: http://dx.doi.org/10.1073/pnas.1416573112

52. Baldo BA, Spencer RC, Sadeghian K, Mena JD. GABA-Mediated Inactivation of Medial Prefrontal and Agranular Insular Cortex in the Rat: Contrasting Effects on Hunger- and Palatability-Driven Feeding. Neuropsychopharmacology 2016; 41: 960–970.

